# Mapping Pathways to Neuronal Atrophy in Healthy, Mid-aged Adults: From Chronic Stress to Systemic Inflammation to Neurodegeneration?

**DOI:** 10.1101/2023.10.18.562886

**Authors:** Julia K Schaefer, Veronika Engert, Sofie L Valk, Tania Singer, Lara MC Puhlmann

**Affiliations:** Cognitive Neuropsychology, Department of Psychology, Ludwig-Maximilians-Universität München; Research Group “Social Stress and Family Health”, Max Planck Institute for Human Cognitive and Brain Sciences, Leipzig, Germany; Institute of Psychosocial Medicine, Psychotherapy and Psychooncology, Jena University Clinic, Friedrich-Schiller University, Jena, Germany; Otto Hahn Group Cognitive Neurogenetics, Max Planck Institute for Human Cognitive and Brain Sciences, Leipzig, Germany; Institute of Neuroscience and Medicine, Brain & Behaviour (INM-7), Research Centre Jülich, FZ Jülich, Jülich, Germany; Institute of Systems Neuroscience, Medical Faculty, Heinrich Heine University Düsseldorf, Düsseldorf, Germany; Social Neuroscience Lab, Max Planck Society, Berlin, Germany; Leibniz Institute for Resilience Research, Mainz, Germany

**Keywords:** chronic stress, low-grade inflammation, hippocampal volume, cortical thickness, structural equation models

## Abstract

Growing evidence implicates systemic inflammation in the loss of structural brain integrity in natural ageing and disorder development. Chronic stress and glucocorticoid exposure can potentiate inflammatory processes and have also been linked to neuronal atrophy, particularly in the hippocampus and the human neocortex. To improve understanding of emerging maladaptive interactions between stress and inflammation, this study examined evidence for glucocorticoid- and inflammation-mediated neurodegeneration in healthy mid-aged adults.

N=169 healthy adults (mean age = 39.4, 64.5% female) were sampled from the general population in the context of the ReSource Project. Stress, inflammation and neuronal atrophy were quantified using physiological indices of chronic stress (hair cortisol and cortisone concentration), systemic inflammation (interleukin-6, high-sensitive C-reactive protein), the systemic inflammation index (SII), hippocampal volume (HCV) and cortical thickness (CT) in regions of interest. Structural equation models were used to examined evidence for pathways from stress and inflammation to neuronal atrophy.

Model fit indices indicated good representation of stress, inflammation, and neurological data through the constructed models (CT model: robust RMSEA = 0.041, robust ***χ***^2^= 910.90; HCV model: robust RMSEA < 0.001, robust ***χ***^2^ = 40.95). We replicated typical negative age-cortical thickness associations (Anterior cingulate cortex (β =-0.51, p < .001), Parahippocampal Cortex (β = −0.50, p = .012), Frontal Lobe (β = −0.56, p < .001) and Temporal Lobe (β = −0.61, p < .001). Among inflammatory indices, only the SII was positively associated with hair cortisol as one indicator of chronic stress (β = 0.18, p<.05). Direct and indirect pathways from chronic stress and systemic inflammation to cortical thickness or hippocampal volume were non-significant.

We identify the SII as a potential marker of systemic inflammation in human psychobiological studies. More generally, these data suggest that neurophysiological associations found in at-risk populations are not detectable in healthy, mid-aged populations. We conclude that inflammation and glucocorticoid-mediated neurodegeneration may only emerge during advanced ageing and disorder processes and may thus have limited use as early risk markers. Future work should examine these pathways in prospective longitudinal designs, for which the present investigation serves as a baseline.

## 1. Introduction

### 1.1. Short Overview

Mental health conditions and other disorders of the brain are highly prevalent and rank among the leading causes for global burden of disease (James et al., 2018; Wittchen, et al., 2011). Chronic stress and pro-inflammatory activity are both linked to neuronal atrophy in cortical and subcortical structures, forming pathways that are implicated in accelerated ageing, cognitive impairment and the development of psychiatric brain disorders, such as Major Depressive Disorder (MDD) (Chrousos, 2009; Chung et al., 2002; Kremen et al., 2010; Lebedeva et al., 2018; Marsland et al., 2015; McEwen, 2008; Sapolsky, 2004). To date, however, few studies have comprehensively investigated the complex interrelation of chronic stress, systemic inflammation, and brain morphology. In particular, little is known about the emergence of their maladaptive interactions and potential pathways to disorder. The present study addresses this gap by comprehensively investigating the interplay between glucocorticoid (GC) exposure, systemic inflammation, and cortical and subcortical brain morphology in a healthy mid-aged sample. Data was collected at baseline of a large-scale, multi-disciplinary longitudinal mental training intervention study, the ReSource Project (Singer et al., 2016). Using structural equation models (SEMs), we evaluate evidence for different neurobiological pathways that may indicate emerging maladaptive processes, which is crucial to identify neurobiological risk factors and targets for future preventive interventions.

### 1.2. Chronic Stress

Among the most important endocrine mediators of the stress response and its long-term health effects are GCs like cortisol, the end-product of the human hypothalamus-pituitary-adrenal (HPA) axis. Released as part of a cascade of stress-mediators, cortisol is an essential signalling agent in mainly down-regulatory feedback loops that centrally involve the brain (McEwen, 2007). Prolonged exposure to stress and GC signalling appears to impair these regulatory mechanisms, potentially via reduced sensitivity to GC signalling (glucocorticoid receptor resistance (GCR) hypothesis, Cohen et al., 2012) leading to a failure to properly terminate HPA axis activity (Chrousos et al., 1993; Chrousos, 1995).

Chronic stress and the resulting sustained GC exposure have been linked to neuronal atrophy and the development of prevalent psychiatric disorders such as MDD (Duman & Monteggia, 2006). Particularly well-documented is the neurotoxic effect of sustained GC exposure in the hippocampus (Geerlings & Gerritsen, 2017; Lupien et al., 1998; McEwen & Gould, 1990; McEwen, 1999; Sapolsky & Pulsinelli, 1985; Sapolsky, 1990), the brain region expressing the highest density of GC receptors (McEwen, 1982). Inverse associations with basal cortisol levels have, however, also been found for regional and total brain volumes (Sigurdsson et al., 2012), and HPA axis dysregulation seems to be linked to smaller left ACC volumes (MacLullich et al., 2006) and frontal lobe atrophy (Gold et al., 2005). Similarly, total diurnal cortisol output is inversely associated with cortical thickness (CT) (Lebedeva et al., 2018). In patients with early-stage MDD, serum cortisol levels were inversely correlated with CT in several brain areas (Liu et al., 2015). Overall, neurotoxic effects of stress and GC exposure thus appear to extend beyond the hippocampus to cortical brain regions (Lupien & Lepage, 2001).

### 1.3. Systemic Inflammation

Similar to the stress response, the acutely adaptive innate immune response can become damaging if not appropriately terminated. Failure to downregulate pro-inflammatory activity can result in systemic inflammation, a maladaptive state that manifests itself with prolonged, low-level elevations of pro-inflammatory cytokines, such as Interleukin-6 (IL-6) and high-sensitive C-reactive Protein (CRP), the most commonly assessed markers of systemic inflammation (Slavich, 2020; Rohleder, 2019). Like chronic stress, systemic inflammation is associated with a range of psychological disorders such as MDD (Rosenblat et al., 2014) and Schizophrenia (Stojanovic et al., 2014). Neuroinflammation and the co-occurrence of systemic inflammation and neuronal atrophy have in particular been implicated in the development of these disorders. Early studies in rats show that neuropathological changes and loss of synapses and granule neurons are associated with chronic neuroinflammation and IL-6 concentrations (Campbell et al., 1993; Heyser et al., 1997; Qiu et al., 1998). IL-6 also appears to modulate neurogenesis in the dentate gyrus of the mouse hippocampus (Vallieres et al., 2002).

In humans, associations between inflammation and brain morphology are commonly studied in clinical samples. Systemic inflammation in terms of elevated CRP, IL-6 and TNF-*α* levels is inversely correlated to lower CT and cortical grey matter volume in patients with schizophrenia (Jacomb et al., 2018; Massuda et al., 2014), and it has been associated with the promotion of neurodegeneration in chronic neurodegenerative diseases, such as Alzheimer’s disease (Holmes et al., 2007).

Similar associations have also been found in subclinical samples, albeit less prominently, providing evidence for an inflammatory pathway towards progressive neuronal atrophy and disorder development. Studies involving healthy subjects report inverse associations between IL-6 or CRP levels and hippocampal grey matter and total brain volume (Gu et al., 2017; Jefferson et al., 2007; Marsland et al., 2008), as well as cortical thinning in middle aged (van Velzen et al., 2017) and elderly individuals without dementia (Fleischman et al., 2010; McCarrey et al., 2014). Biological ageing processes are accompanied by enhanced levels of inflammatory markers (Godbout & Johnson, 2004; Wei et al., 1992; Ye & Johnson, 1999;) and also appear to play an important role in the interplay of chronic stress and systemic inflammation (Gouin et al., 2008). Thus, early onset of inflammation-mediated neuronal atrophy may serve as a risk marker for accelerated ageing and neurodegenerative disorders.

### 1.4. Stress, Inflammation, and Brain Structure

Chronic stress and cortisol exposure closely interact with systemic (or chronic low-grade) inflammation. While GCs generally have a regulatory effect on the acute immune response (Waage et al., 1990), prolonged psychosocial stress is associated with elevated low-grade inflammation (Rohleder, 2014, 2019). It is thus presumed that chronic stress may alter GC signalling and lead to a pro-inflammatory effect (Ader et al., 1995; Arimura et al., 1994; Black, 2002; Chrousos, 2000; Hänsel et al., 2010; McEwen et al., 1997). The GC receptor hypothesis for example assumes that due to permanent exposure to GCs, not only receptors in hypothalamus and pituitary but also in immune cells such as macrophages become insensitive to GCs, which can lead to the disruption of GC-induced suppression of inflammation (Cohen et al., 2012; Miller et al., 2008; Stark et al., 2001). Multiple human studies suggest a link between increased stress experience and inflammation, including in healthy adults (Maes et al., 1998; Miller et al., 2002). Chronic stress and systemic inflammation are highly synergistic in their interactive effect on many pathologies such as Metabolic Syndrome (MtS) (Almadi et al., 2013) or coronary artery disease (Nijm & Jonasson, 2009). As mentioned above, MDD was found to be closely related to inflammatory processes, but it also interacts with stress in inhibiting the negative feedback loop of inflammation, adding to the enduring state of inflammation, especially pronounced in aged subjects (Robles et al., 2016).

Although the interplay between chronic stress and systemic inflammation and their joint contribution to alterations in brain morphology has been subject to several high-profile reviews, studies examining these associations in a joint statistical model and in a healthy sample are rare. Summarizing the animal literature, Sorrells and Sapolsky (2007) and Kubera et al. (2011) conclude that in animal models, stress-induced inflammation enhances neurodegeneration, which in turn may provoke depression-like behaviours (see also inflammatory and neurodegenerative hypothesis, Maes et al., 2009). Fewer studies have been able to investigate this maladaptive triangulation in humans, although one review on MDD patients identifies similar relations on chronic stress, neuroinflammation and alterations in brain structure and function (Kim & Won, 2017). Regarding endocrine stress markers, reduced GC responsiveness and enhanced IL-6 levels were also related to thinner cortices in patients with mood disorders (van Velzen et al., 2017) and to smaller hippocampi for patients with MDD specifically (Frodl et al., 2012).

### 1.5. Present Study

Beyond the above clinical studies, little is known about the relation between chronic stress, systemic inflammation, and brain structure in healthy adults and the general population. As such, it remains unclear to what extent chronic stress and systemic inflammation are linked to neuronal atrophy in the absence of disorder or advanced ageing, and whether stress and inflammation interact or potentiate each other as risk factors for neurodegenerative processes. The present work examines this question, aiming to inform neurobiological models of disorder development and chronic stress and inflammation as risk indices for early neurodegenerative processes. To map the interrelation of physiological indices related to chronic stress, systemic low-grade inflammation, and cortical and subcortical brain morphology, we used multimodal cross-sectional data from N=169 healthy adults (N=150 for subcortical morphology). Data collected at baseline of a large-scale, multi-disciplinary longitudinal mental training intervention study, the ReSource Project (Singer et al., 2016). Chronic stress was measured via hair cortisol (HCC) and cortisone (HEC) concentrations. Systemic inflammation was indicated by blood serum levels of IL-6, hs-CRP and the systemic inflammation index (SII). Finally, we examined brain morphology via hippocampal volume (HCV), since hippocampal structure and function are closely tied to stress and neuroinflammation, as well as via thickness of the neocortex (cortical thickness, CT). CT provides an anatomically specific (Lemaitre et al., 2012; Winkler et al., 2010) and particularly sensitive measures of grey matter variation, especially in ageing (Hutton et al., 2009), for example compared to volume-based methods.

In previous work of the ReSource Project, we demonstrated the multidimensionality of the psychophysiological construct stress and its relation to various health and sleep measures using network analysis (Engert et al., 2018). Here, we now examine inflammation and stress as latent constructs and in their relation to brain morphology. Using SEMs, we test secondary hypotheses on specific physiological pathways to neurostructural atrophy involving mediation and moderation effects through stress and inflammation: We expected a positive association between the latent constructs chronic stress and systemic inflammation, representing stress-related inflammation, potentially mediated via the BMI, which we previously found associated with single inflammatory and stress-related biomarkers in the same sample (Engert et al., 2018). We also expected a cumulative negative effect of elevated chronic stress and systemic inflammation on both CT and HCV, in form of either an indirect effect of stress effects via inflammation, or a moderation effect of inflammation and stress. Finally, next to IL-6 and hs-CRP as our primary indicators of systemic inflammation, we further tested an indirect association from chronic stress to brain structure via the systemic inflammation index (SII) which is assumed to have prognostic value for overall survival in certain cancers (Hong et al., 2015; Zhong et al., 2017) but has not yet been examined in humans with regard to psychosocial factors such as stress-related inflammation.

## 2. Methods

### 2.1. Sample and Recruitment

Data for the present investigation was collected in the context of a large-scale 9-month longitudinal mental training study, the ReSource Project (Singer et al., 2016). Healthy participants with an age range of 20 – 55 years (mean age = 39.4, SD = 9.8) were recruited (see Tables 2a, 2b). All participants underwent mental and physical health screenings as well as two clinical diagnostic interviews [Structured Clinical Interview for DSM-IV Axis-I (SCID-I) (Wittchen & Pfister, 1997); SCID-II for Axis-II disorders (First et al., 1997)]. Participants were excluded if they fulfilled the criteria for an Axis I disorder in the past two years or an Axis-II disorder at any time in their life. Additional inclusion criteria were several chronic physical pathologies and intake of medication affecting the HPA axis or central nervous system. A detailed description of the recruitment procedure and information about the final sample of the ReSource Project can be found in Singer et al., 2016, chapter 7. The ReSource Project was registered via the Protocol Registration System of ClinicalTrial.gov (Identifier NCT01833104) and the study was approved by the research ethics boards of Leipzig University (ethic number: 376/12-ff) and Humboldt University Berlin (ethic numbers: 2013–20, 2013–29, 2014–10). Participants gave written informed consent, received financial compensation, and could withdraw from the study at any time.

For the present investigation, only data collected at the pre-training baseline (T0) of the ReSource Project was evaluated. Although the data reported here were previously published in the context of other research questions mostly pertaining to the effect of ReSource training (Degering et al., 2023; Engert et al., 2018; Puhlmann, Engert, et al., 2019; Puhlmann, Linz, et al., 2021; Puhlmann, Valk, et al., 2019; Puhlmann, Vrtička, et al., 2021; Valk et al., 2017, 2023), none of these studies investigated the complex relation between measures of chronic stress physiology, inflammatory activity and brain morphology, and potentially associated pathways of moderation and mediation. The present study is an a-posteriori exploratory study not planned during the designing of the ReSource Project and all formulated hypotheses and models should be considered secondary.

### 2.2. Measures

#### 2.2.1. Indices of Chronic Stress: Hair cortisol (HCC) and Hair Cortisone Concentration (HEC)

A popular biomarker of chronic stress is the extraction of HCC, and HEC as a complementary measure, which both serve as indices of systemic cortisol exposure (Short et al., 2016; Stalder & Kirschbaum, 2012; Stalder et al., 2012). HCC appears to be quite robust to confounders and is associated with well-known correlates of stress-related cardiometabolic parameters such as systolic blood pressure and BMI (Stalder et al., 2017). Both HCC and HEC are generally more stable compared to serum or salvia cortisol levels that are part of a dynamic system with day-to-day changes in activity (Ross et al., 2014). For their assessment, hair strands were collected close to the scalp and a 3 cm segment, corresponding to approximately 3 months of cortisol exposure, was analysed. Concentrations of HCC and cortisone were measured with liquid chromatography-tandem mass spectrometry (LC–MS/MS) (Gao et al., 2016).

#### 2.2.2. Indices of Systemic Inflammation: Interleukin-6 (IL-6) and high-sensitive C-Reactive Protein (hs-CRP)

IL-6 and hs-CRP were used as primary indices of systemic inflammation. For the assessment of IL-6 and hs-CRP levels, 5.5 ml blood was collected and stored at −80 degrees Celsius. Hs-CRP was measured with a latex-enhanced immunoturbidimetric assay using the Siemens Advia 1800 Clinical Chemistry System (Siemens Healthineers, Tarrytown, NY, USA). IL-6 levels were detected with a solid phase enzyme-labelled chemiluminescence immunometric assay using the random access chemiluminescence-immunoassay system (IMMULITE 2000, Siemens Healthineers, Tarrytown, NY, USA) (for more details see Engert et al. (2018)). Levels of IL-6 follow a circadian cycle, with lower levels during daytime and higher levels during the night (Vgontzas et al., 2005). To account for these fluctuations, time of sampling was documented and included as a control variable in all analysis.

#### 2.2.3. Systemic Inflammation Index (SII)

Systemic inflammation is a complex and extensive process, during which not only levels of IL-6 and hs-CRP but also the count of circulating leukocytes such as neutrophile granulocytes and monocytes is increased while the lymphocyte count is decreased (Rink et al., 2015). Some studies use ratios of neutrophils, thrombocytes (platelets) and lymphocytes as indicators for systemic inflammation (Systemic inflammation Index, SII; Hong et al., 2015; Wu et al., 2016). The SII can thus be derived from a complete blood count and is calculated as the product of thrombocytes and neutrophils divided by lymphocytes (Hong et al., 2015; Hu et al., 2014; Wu et al., 2016).

The SII is assumed to have prognostic value for overall survival in certain cancers (Hong et al., 2015; Zhong et al., 2017). Even though the SII is an index of systemic inflammation, it has not yet been examined with regard to psychosocial factors in humans. In the current study, IL-6 and hs-CRP were considered as the main marker of systemic inflammation, and the SII was related to chronic stress and brain structure in an additional analysis.

#### 2.2.4. MRI Acquisition

High resolution T1-weighted structural MRI images were acquired on a 3T Trio TIM scanner (Siemens Verio; Siemens, Erlangen, Germany) with a 32 - channel head coil, using magnetization-prepared rapid gradient echo (MPRAGE; 176 sagittal slices; repetition time, 2300 milliseconds; echo time, 2.98 milliseconds; inversion time, 900 milliseconds; flip angle, 7; field of view, 240256 mm2; and matrix, 240256; 111 mm3 voxels) sequence.

#### 2.2.5. Cortical Thickness (CT) Calculation and selection of ROIs

We used Freesurfer version 5.1.0 (consistent with previous publications from the ReSource Project, e.g. Valk et al., 2017) to generate cortical surface models for the calculation of CT following previously reported steps (Dale et al., 1999; Fischl et al., 1999; see also Valk et al., 2017). Briefly, T1-weighted images were intensity normalized and skull stripped, and the grey/ white matter cortical boundary tessellated. After automatic correction of topology, the surface deformations converged the cortical interfaces of the inner boundary (gray/white matter) and outer boundary (gray matter/ cerebrospinal fluid), following intensity gradients. Surface reconstruction was visually inspected by two independent raters and inaccuracies manually corrected. CT was then calculated as the shortest distance from the gray/white matter boundary to the gray matter/CSF boundary at each vertex on the tessellated surface. For more details of the processing steps see Dale et al., (1999); Fischl et al., (1999) and Han et al., (2006). Regions of interest (ROIs) for CT analyses were parcellated following the Desikan-Killiany Atlas as implemented in FreeSurfer 5.1.0.

To not overload our models, and since it is advised to build SEMs on a strong conceptual foundation (Bentler & Chou, 1987; Hoyle, 1995), we focused on CT of 14 regions of interest identified from the literature. We compared ROIs of studies with healthy samples (Kaur et al., 2015; Kremen et al., 2010; Marsland et al., 2015; Piras et al., 2012; Savic, 2015; van Velzen et al., 2017), pathological samples (Chiappelli et al., 2017; Jacomb et al., 2018; Lebedeva et al., 2018; Liu et al., 2015; Massuda et al., 2014; Ottino-Gonzalez et al., 2017; Veit et al., 2014;), aged samples (Fleischman et al., 2010), longitudinal studies (Gu et al., 2017; McCarrey et al., 2014) and two reviews (Byrne et al., 2016; Sheline, 2003), selecting ROIs in which CT had been found to relate to either HCC/HEC or IL-6/CRP. All ROIs and the final factor solution for ROIs is presented in Table 3 (for more details on ROI selection see Supplementary Methods C).

#### 2.2.6. Hippocampal Volume (HCV) Calculation

On the base of the high resolution T1-weighted structural MRI images CA1-3, CA4/DG, and subiculum (SUB) were segmented, with a patch-based algorithm in every subject. Briefly, the algorithm employs a population-based patch normalization relative to a template library (Kulaga-Yoskovitz et al., 2015), which has shown high segmentation accuracy of hippocampal subfields in previous validations (Caldairou et al., 2016). All HCV segmentations were quality controlled by two independent raters and any segmentations with average quality rating scores lower than 5 were excluded from the analysis. (details on the algorithm and quality control procedure see Puhlmann, et al., 2021). While Freesurfer also provides estimates of HCV, we use the patch-based method throughout the ReSource Project following our preregistered study. The resulting surface-based estimates show decent overlap with Freesurfer estimates (Puhlmann et al., 2021).

#### 2.2.7. Other Measures

The body mass index (BMI), as the relation of the individual’s body weight in kilograms to the squared height, was incorporated as an indicator for adipose tissue.

### 2.3. Data Analysis

Data analysis was conducted using Structural Equation Models (SEMs). SEMs allow the testing of complex interrelations by representing conceptual research models through a system of connected regression-style equations. An additional benefit of SEMs is the possibility to include latent factors, which are estimated based on multiple indicator variables via factor analysis. In our hypothesized model, we indicated chronic stress via HCC and HEC, systemic inflammation via IL-6 and hs-CRP, and CT via the selected ROIs, based on the above reviewed evidence. Using multiple indices increases the reliability of latent factors and reduces the influence of random measurement noise. Total left and right HCV were added as measurement variables without forming a latent construct as did literature did not indicate subfield specific associations.

#### 2.3.1. Sample Size Calculation

Following recommendations to ensure adequately powered SEMs (MacCallum et al., 1996; Westland, 2010), we assessed whether the pre-existing sample size was sufficient for the planned model using the *A-priori Sample Size Calculator for Structural Equation Models* (Soper, 2022). Given the levels of complexity in both models, the available sample sizes of N= 169 respectively N = 150 could be considered sufficient. For more details on the sample size calculation see *Supplementary Methods E*.

#### 2.3.2. Variable pre-processing

The biological variables IL-6, hs-CRP, HCC and HEC were ln-transformed to remedy their typical skewed distribution. Outliers defined as *+/-SD* = 3 were winsorized to the upper or lower boundary of 3 SDs, respectively. For more details on the statistical pre-processing of variables see *Supplementary Methods D*.

#### 2.3.3. Fitting the SEMs

To address our conceptual model of interrelations, we fit one SEM to map the chronic stress and systemic inflammation in relation to CT, and one in relation to HCV (Figure 1 and 2, respectively). Chronic stress and systemic low-grade inflammation were included as latent factors as described above, with one indicator variable fixed to *λ* = 1, as recommended for hypothesis-driven measurement models with few indicator variables (Hayduk & Littvay, 2012). The SII was included exploratorily as an additional endogenous variable. BMI was modelled as a mediator from chronic stress to systemic inflammation following previous results (Engert et al., 2018). Age, hormonal status (male, female no cycle, female hormonal contraceptives, female natural cycle) and information about smokers/ non-smokers were always included as exogenous variables (i.e., variables that perform only as independent variable) to account for their well-established influence on cortisol/cortisone, inflammatory proteins and brain structure (Fleischman et al., 2010; Godbout & Johnson, 2004; Kajantie & Phillips, 2006; Thayer et al., 2010; Ugur et al., 2018; Veit, R., et al., 2014; Wright et al., 2006).

**Figure 1:**
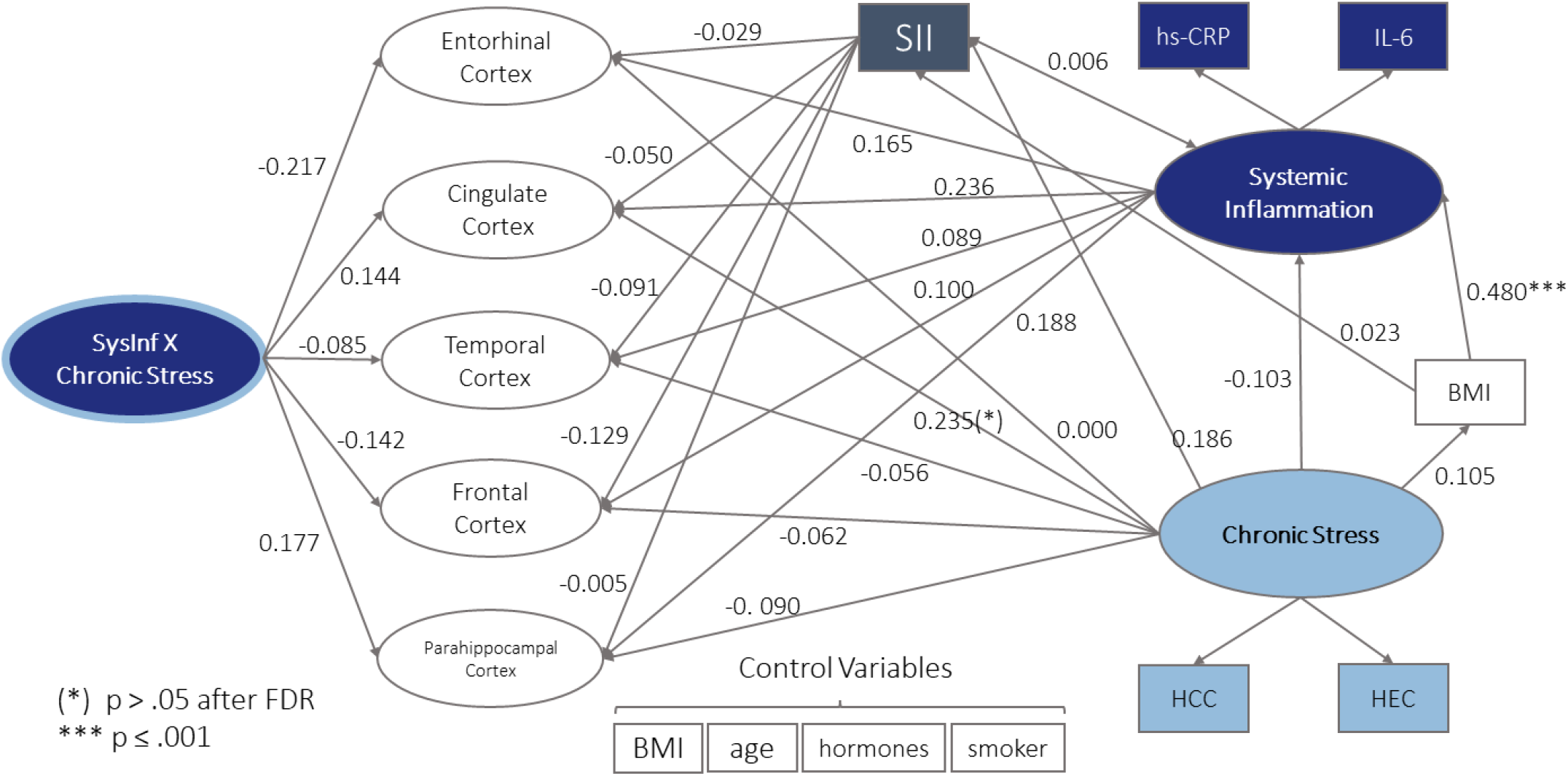
Structural model with all latent factors of CT, latent factor chronic stress with indicator variables HCC and HEC, latent factor systemic inflammation with indicator variables IL-6 and hs-CRP and interaction factor of inflammation and chronic stress. Standardized (latent and observed variables) path coefficients are reported. All variances, indicator variables for the CT ROIs and covariations of error terms are hidden for visual clarity. Spheres represent latent factors, square boxes measured variables. BMI is modelled as a control variable for all variables except for chronic stress and systemic inflammation, where it was modelled as a mediator variable.

**Figure 2:**
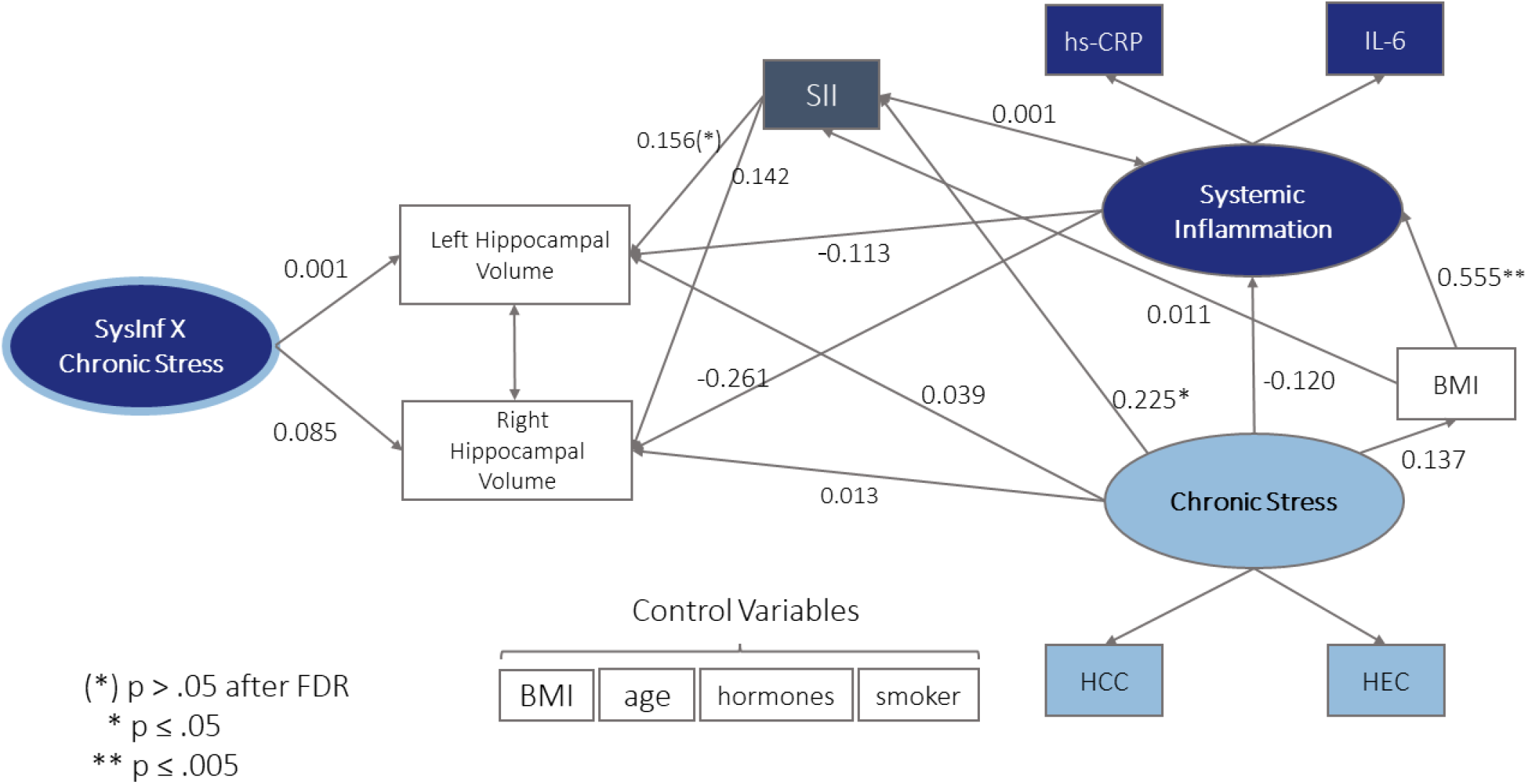
Structural model with HCV in left and right hemisphere, latent factor chronic stress with indicator variables HCC and HEC, latent factor systemic inflammation with indicator variables IL-6 and hs-CRP and interaction factor of inflammation and chronic stress. Standardized (latent and observed variables) path coefficients are reported. All variances, indicator variables for the CT ROIs and covariations of error terms are hidden for visual clarity. Spheres represent latent factors, square boxes measured variables. BMI is modelled as a control variable for all variables except for chronic stress and systemic inflammation, where it was modelled as a mediator variable.

As the first physiological endpoint, CT was added to the SEM. To robustly represent CT without averaging across functionally and structurally heterogeneous regions, we formed five latent CT factors based on the ROI estimates (Table 1). For more details on the formation of latent factors of cortical thickness see *Supplementary Methods C*.

**Table 1.** Latent factor solution of ROIs.

| Latent Variable | ACC | Frontal Lobe | Temporal Lobe | Entorhinal Cortex | Parahippocampal Cortex |
| --- | --- | --- | --- | --- | --- |
| Indicator Variables | left Frontal Rostral ACC | left Frontal Superior Gyrus | left Temporal Fusiform Gyrus | left Entorhinal Cortex | left Parahippocampal Cortex |
|  | right Frontal rostral ACC | right Frontal Superior Gyrus | right Temporal Fusiform Gyrus | right Entorhinal Cortex | right Parahippocampal Cortex |
|  | left Frontal Caudal ACC | left Frontal Caudal Middle Gyrus | left Superior Temporal Banks |  |  |
|  | right Frontal Caudal ACC | right Frontal Caudal Middle Gyrus | right Superior Temporal Banks |  |  |
|  |  | left Paracentral Gyrus | left Temporal Inferior Gyrus |  |  |
|  |  | right Paracentral Gyrus | right Temporal Inferior Gyrus |  |  |
|  |  | left Precentral Gyrus | left Transverse-temporal Gyrus |  |  |
|  |  | right Precentral Gyrus | right Transverse-temporal Gyrus |  |  |
|  |  |  | left Temporal Middle Gyrus |  |  |
|  |  |  | right Temporal Middle Gyrus |  |  |
|  |  |  | left Temporal Superior Gyrus |  |  |
|  |  |  | right Temporal Superior Gyrus |  |  |

Final five latent factor solution, each latent factor listed with all its ROI indicator variables.

The second model relating chronic stress and systemic inflammation to HCV was identical to the model comprising CT, except that all latent factors of CT were replaced with the two exogenous variables HCV in the left and right hemisphere.

#### 2.3.4. Path analyses and model comparisons

To test the statistical significance of direct associations, potential moderation effects and indirect associations between the latent constructs and indicator variables of interest, we conducted path-analyses within the two fitted SEMs. Indirect associations were evaluated within an implicit procedure (Rungtusanatham et al., 2014), testing for the joint significance of every constituent path of an indirect association. For moderation analysis, product indicators for latent interaction factors were calculated, following the residual centring approach (Little et al., 2006), which is also recommended by Steinmetz et al. (2011). All path coefficient estimates are reported in the all-variables-standardized-version, *Std.all.* For evaluating statistical significance an *α*-level of .05 was applied. Family-wise error correction was performed on significant parameters by applying the false discovery rate (FDR) (Benjamini & Hochberg, 1995) to correct for multiple comparisons of paths to each of the different brain areas included in the model. Once all models were set, direct model comparisons of nested models were evaluated through significance testing of chi^2 differences.

## 3. Results

### 3.1. Final Sample

From the *N* = 332 subjects included at study baseline (T0) (Singer et al., 2016), *n* = 169 provided data for all present variables of interest in the CT model and *n* = 150 in the HCV model and could thus be used in the SEM analysis (see Table 2a) and b), for more details see also *Supplementary Table S1*).

**Table 2a.** Sample Characteristics CT for model (N=169)

|  | Female | male | overall |
| --- | --- | --- | --- |
|  | (N=109) | (N=60) | (N=169) |
| Mean age (SD) | 41.0 (9.40) | 36.6 (9.84) | 39.4 (9.75) |
| Mean BMI (SD) | 23.0 (3.29) | 24.3 (2.84) | 23.5 (3.19) |
| Median SII (Gpt/l) [range] | 471 [170, 1280] | 406 [147, 1320] | 445 [147, 1320] |
| Median IL-6 (pg/ml) [range] | 1.49 [1.28, 24.6] | 1.44 [1.28, 3.40] | 1.47 [1.28, 24.6] |
| Median hs-CRP (mg/L) [range] | 0.925 [0.128, 13.2] | 0.500 [0.138, 5.98] | 0.709 [0.128, 13.2] |
| Median HCC (pg/mg) [range] | 3.31 [0.486, 95.2] | 4.81 [0.181, 52.1] | 3.83 [0.181, 95.2] |
| Median HEC (pg/mg) [range] | 8.89 [1.87, 66.1] | 14.6 [2.54, 51.0] | 11.0 [1.87, 66.1] |

**Table 2b.** Sample Characteristics for HCV model (N=150)

|  | Female | male | overall |
| --- | --- | --- | --- |
|  | (N=98) | (N=52) | (N=150) |
| Mean age (SD) | 40.9 (9.52) | 36.1 (9.34) | 39.2 (9.70) |
| Mean BMI (SD) | 23.1 (3.41) | 24.4 (2.91) | 23.6 (3.30) |
| Median SII (Gpt/l) [range] | 477 [170, 1280] | 406 [178, 1320] | 452 [170, 1320] |
| Median IL-6 (pg/mL) [range] | 1.49 [1.29, 24.6] | 1.44 [1.28, 2.10] | 1.48 [1.28, 24.6] |
| Median hs-CRP (mg/L [range] | 0.943 [0.128, 13.2] | 0.532 [0.138, 5.98] | 0.741 [0.128, 13.2] |
| Median HCC (pg/mg) [range] | 3.59 [0.486, 95.2] | 4.63 [0.181, 52.1] | 3.99 [0.181, 95.2] |
| Median HEC (pg/mg) [range] | 8.85 [1.87, 61.8] | 13.7 [2.54, 45.6] | 10.9 [1.87, 61.8] |

Missing data was excluded case wise, as implemented by the lavaan (Rosseel, 2012) and sem (Fox, 2006) packages to ensure a true and unbiased correlation matrix as input for the SEM. Most cases were excluded due to missing HCC or HEC data, because sampling of hair for the assessment of HCC and HEC was presented to participants as an optional rather than a core testing procedure, leading to lower adherence rates (see Puhlmann et al., 2021 for further details).

### 3.2. Correlations of stress and inflammation biomarkers

Before building latent constructs, partial correlations between the key risk factors in our hypothesized pathways were calculated, namely the chronic stress indicator variables HEC and HCC, inflammation indicators hs-CRP and IL-6, as well as SII and BMI. We replicated previously identified associations (see Engert et al., 2018) and additionally found that the SII was significantly positively correlated with HCC (p<.05) (see Table 3).

**Table 3.**
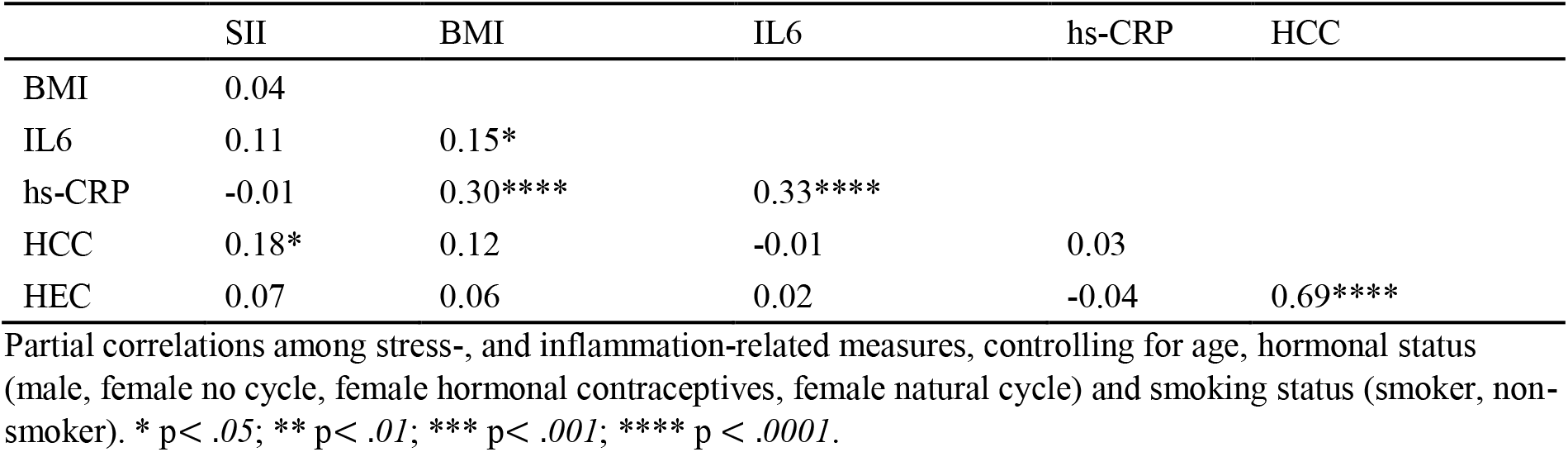
Partial Correlations among stress-, and inflammation-related measures.

|  | SII | BMI | IL6 | hs-CRP | HCC |
| --- | --- | --- | --- | --- | --- |
| BMI | 0.04 |  |  |  |  |
| IL6 | 0.11 | 0.15* |  |  |  |
| hs-CRP | -0.01 | 0.30**** | 0.33**** |  |  |
| HCC | 0.18* | 0.12 | -0.01 | 0.03 |  |
| HEC | 0.07 | 0.06 | 0.02 | -0.04 | 0.69**** |

Partial correlations among stress-, and inflammation-related measures, controlling for age, hormonal status (male, female no cycle, female hormonal contraceptives, female natural cycle) and smoking status (smoker, non-smoker). \* $p < .05$ ; \*\* $p < .01$ ; \*\*\* $p < .001$ ; \*\*\*\* $p < .0001$ .

In initial sanity checks, we also confirmed the significance of several common associations not directly related to our conceptual research model, such as negative associations of age with latent CT factors and HCV, and positive associations between BMI and inflammation (see *Supplementary Tables S8 and S9*).

### 3.3. Cortical Thickness Model

Setting up the full CT Model (N=169) as described above (see Figure 1), resulted in an overidentified model with good model fit indicated by most model fit measures (robust chi^2 (910.904) < 2*df (706), robust CFI (.929), robust TLI (.918), robust RMSEA (.041), robust SRMR (.059)). Path analysis indicated that the factor chronic stress was not associated with any factor representing CT (see *Supplementary Table S2)*. Although there was a significant effect of chronic stress on CT in the anterior cingulate cortex, this effect was no longer significant after correcting for multiple comparisons with the positive false discovery rate (see Figure 1). Similarly, there was no significant indirect association of chronic stress and CT via systemic inflammation (Figure 1).

Path analysis further showed that systemic inflammation was not associated with any factor representing CT (see Figure 1) and that chronic stress did not play a moderating role in the association of systemic inflammation and CT (see *Supplementary Table S3)* and Figure 1).

### 3.4. Hippocampal Model

Setting up the HCV model, modelling the same paths as in the CT model, two Heywood cases occurred. They were handled by setting the product indicator variables to be equal (for more details on the handling of Heywood cases see *Supplementary Methods B*). All model fit indices and parameter estimates in the HCV model are reported in the Heywood case corrected version. Thus, the full HCV model (see Figure 2), too, resulted in an overidentified model with good model fit (robust chi^2 (40.945) < 2*df (60), robust CFI (1.000), robust TLI (1.117 truncated to 1.000) robust RMSEA (.000), robust SRMR (.060)) (see Hu & Bentler, 1999; MacCallum et al., 1996).

For the Hippocampal Model, similar to CT, path analysis showed no associations of either chronic stress or systemic inflammation with the left or right HCV (see Figure 2; *Supplementary Table S4 and S5)*). There was also no significant specific indirect association with chronic stress via systemic inflammation and chronic stress did not play a moderating role in the relation between systemic inflammation and HCV.

All results of the path analyses were confirmed when addressing the same hypothesized paths via model comparison of constrained models with the paths of interest individually fixed to zero, compared to unconstrained models.

### 3.5. Exploratory analysis

In exploratory path analyses, the association of SII with factors of CT and HCV were evaluated, as well as an indirect association of chronic stress via the SII (see Figure 1 and 2). None of these associations was significant in the CT model (see *Schaeferementary Table S6*). In the HCV model the SII was significantly positively associated with the left HCV (see *Supplementary Table S7*), this effect was no longer significant after correcting for multiple comparisons with the positive false discovery rate. The SII was furthermore significantly related to the latent factor chronic stress (*p<.05*) in the HCV model (see Table 3 and Figure 2).

## 4. Discussion

Chronic stress and related glucocorticoid (GC) exposure are linked to systemic inflammation (Chrousos, 2000; Cohen et al., 2012; Hänsel et al., 2010), and both processes have been implicated in advanced neurodegeneration (Gu et al., 2017; Jefferson et al., 2007; Kim & Won, 2017; Lebedeva et al., 2018; Lupien et al., 1998; Marsland et al., 2008; McEwen & Gould, 1990; McEwen, 1999). Less is known, however, about the relation of biomarkers of low-grade inflammation, chronic stress and brain morphology in healthy subclinical populations. The present study adopted a structural equation modelling (SEM) approach to map these relations in a population-based sample of healthy adults, recruited in the context of the ReSource Project (Singer et al., 2016), including the influence of age and BMI, with the aim of informing the use of these indices in future preventive healthcare approaches.

Models replicated patterns of associations between age and cortical thickness (Salat et al., 2004), age and BMI, and sex and BMI (Heymsfield et al., 1993; Mazariegos et al., 1994). In line with other studies (Wright et al., 2006) we also find a positive association of the latent systemic inflammation factor with the body mass index (BMI). This result replicates our earlier work in the same participants, showing a link between BMI and specifically IL-6 levels in a network analysis investigating the multidimensional interrelations of a large set of stress- and health-related measures (Engert et al., 2018). Subcutaneous adipose tissue is a contributor to increased levels of cytokines and especially IL-6 (Kern et al., 2001; Mohamed-Ali et al., 1997), properties that seem to be represented well in our latent inflammation factor. However, none of the formulated expectations could be supported in this sample. Chronic stress was not associated with HCV or any CT in the identified ROIs, directly or indirectly via systemic inflammation. Similarly, systemic inflammation, was neither directly associated with HCV or CT, nor was this association moderated by chronic stress. The systemic inflammation index (SII) based on neutrophil, thrombocyte and lymphocyte cell counts emerged as a potentially interesting additional inflammatory marker that was associated with HCC, although this link was rendered nonsignificant when HCC and HEC were grouped into a latent chronic stress factor.

Especially in patients and at-risk groups such as older adults, evidence for a link between neuronal atrophy and chronic enhanced cortisol levels (Lebedeva et al., 2018) as well as systemic inflammation (Fleischman et al., 2010; Jefferson et al., 2007; Kaur et al., 2015; van Velzen et al., 2017) is substantial. Chronic stress and systemic inflammation have also been quite reliably associated (Arimura et al., 1994; Black, 2002; Chrousos, 2000; Cohen et al., 2012; Hänsel et al., 2010; McEwen et al., 1997; Munck & Náray-Fejes-Tóth, 1994; Stark et al., 2001). Thus, it is likely that the absence of associations between chronic stress, systemic inflammation and brain structure in the present study are related to the sample demographics. Participants were thoroughly screened for health at the beginning of the ReSource project as it was a 9-month intense longitudinal training study (Singer et al., 2016) and even for a healthy sample displayed relatively low inflammatory levels of CRP (comparing the current sample medians to serum levels considered normal, CRP (mg/L) current median = .709; ref median = 2.8; see Table 2 and Ridker et al., 2000). Accordingly, the results suggest that it is likely that maladaptive interactions only become pronounced as degenerative processes begin to take hold. This may prompt the conclusion that preventive interventions should best be focused on these sensitive periods and at-risk samples. In our own previous work in this sample, we also found that a meditation-based mental training with potential health benefits only reduced CRP and IL-6 values of participants with elevated levels at baseline (Puhlmann et al., 2019).

To map transitions from health to disorder, future studies at this intersection should attempt to identify critical levels of GCs and cytokines for risk and degenerative processes, already in sub-clinical samples.

Many of the associations between chronic stress, systemic inflammation and atrophy of brain structure are strongly influenced by age and more pronounced in elderly subjects (Buford & Willoughby, 2008; Chung et al., 2002; Godbout & Johnson, 2004; Gouin et al., 2008; Kiecolt-Glaser et al., 2003; Licastro et al., 2005; Marsland et al., 2015; Weaver et al., 2002; Wei et al., 1992; Ye & Johnson, 1999;), where they emerge even in the absence of disorder (e.g., Gu et al., 2017). With an age range of 20-55 years and a mean age of 40.7 years, the current mid-aged sample was younger than the samples of older to old adults commonly examined in studies that find associations between stress, inflammation and CT (e.g., mean ages of 55, 59, 69 and 81; Fleischman et al., 2010; Kremen et al., 2010; Lebedeva et al., 2018; McCarrey et al., 2014). It can be extrapolated that effects emerge only as ageing proceeds. Future work should address this research question using samples with even broader age ranges that include old and very old individuals and, ideally, in prospective longitudinal studies.

Although stress and inflammation have been found to affect brain structure in many studies (DePablos et al., 2006; Kubera et al., 2011; Ottino-Gonzalez et al., 2017; Sorrells & Sapolsky, 2007), there is still no consensus on the scope, characteristics and direction of this effect. Possibly, other pathways than hypothesized here may converge better in healthy samples, especially when combined with more nuanced measurement approaches. For example, chronically enhanced levels of GCs can potentially have different, even opposing effects in the central nervous system and the periphery, and the precise neurotoxic effect of GC-related inflammation also depends on the specific brain area of inflammation (Sorrells & Sapolsky, 2007). SEMs can be employed to test a combination of pro- and anti-inflammatory GC pathways in future investigations.

Rather than adopting a more nuanced approach, as we did in this study, other studies have opted to analyse the combined burden of chronic stress and inflammation via the allostatic load (AL) index. This integrative measure of prolonged stress exposure and associated physiological sequelae, including inflammation and metabolic changes, has been identified as a correlate of cortical structure (Juster et al., 2010; McEwen, 1993; Ottino-Gonzalez et al., 2017). While this might be a promising approach in terms of identifying at-risk groups and monitoring overall health trajectories, we argue that more nuanced systemic models are necessary to understand the emergence of disorder and potential therapeutic pathways, such as stress-reduction, more fully.

Relatedly, correlations of individual biomarkers showed that the SII, but not IL-6 or hs-CRP, was significantly associated with HCC. This is one indication that the SII may be a valuable contribution to the construct of systemic inflammation when it comes to associations with physiological chronic stress. The lack of correlation with IL-6 and hs-CRP confirms that it captures divergent aspects of inflammation and underscores that some associations are only revealed in granular approaches that differentiate distinct markers of the same construct.

### 4.1. Strength and weaknesses

Previous studies of chronic stress, systemic inflammation and neuronal atrophy have mostly examined only two out of the three variables at a time. Here, we took a more comprehensive approach and jointly modelled all three variables, allowing us to examine multiple pathways of association simultaneously, while also considering the risk factors age, BMI, hormonal status and smoking status. By conducting a sample size calculation, we ensured our sample size to be sufficient for the estimation of both our models. Although this approach is a significant strength of our study, issues with variable convergence to latent factors may also have hindered pathway detection. Notably, hs-CRP and IL-6 did not show high shared variance in their formative latent factor. This raises the question of whether IL-6 and hs-CRP share sufficient variance to be grouped, or if they should rather be considered as separate measures of systemic inflammation. Furthermore, although two indicator variables for a latent construct are sufficient in some cases (Hayduk & Littvay, 2012), three or more indicator variables are often recommended (Bentler & Chou, 1987; Hayduk & Littvay, 2012).

In general, SEMs and the hypothesized pathways might converge better in samples with naturally higher variation in stress and inflammation markers, such as ageing and patient populations.

Another strength of the present study is the selection of literature-based regions of interest (ROIs) for CT as well as the literature-based assumptions about the modelled paths. This approach is, however, also relatively conservative, working only with CT averages in previously identified regions. Whole-brain exploratory analyses with single indicator variables like CRP or IL-6 may be more sensitive to subtle associations, but do not allow the mapping of complex pathways.

Finally, while the present investigation is informative for preventive healthcare approaches, future studies may focus on more diverse samples or sensitive periods. Next to ageing, consequences of a permanent exposure to stress are also particularly severe in children, especially if chronic stress is experienced during the developmental period (Björntorp, 2001; Pervanidou & Chrousos, 2012). Pathway analyses in the context of longitudinal studies will be crucial to establish a quasi-causal chain between chronic stress, systemic inflammation and neurodegeneration in humans.

### 4.2. Conclusion

A better understanding of the interplay between chronic stress and systemic inflammation in their common contribution to neurodegeneration is crucial to combat stress-related disorders that emerge from cumulative burden in this interdependent system (McEwen, 2000, 2007). The present study used SEMs for nuanced modelling of the relation between chronic stress, systemic inflammation and brain morphology as latent constructs. Models identified no evidence for meaningful associations between these three constructs in a sample of healthy adults from the general population. We conclude that neurophysiological associations found previously in at-risk populations of either much older or diseased participants may not be detectable in the absence of such vulnerability. This suggests that other indices may be more informative as early markers of risk and vulnerability for neurodegenerative disorders. Although latent constructs did not covary as expected, multivariate models were successfully fit and replicated established associations for example between age and neuronal atrophy. These findings can serve as a baseline for studies investigating similar research questions in pathological or ageing populations. We further identified the SII as a potential informative marker of systemic inflammation in human psychobiological studies. Overall, we advocate the use of both the SII and path modelling in future studies to do justice to the complexity and interconnectivity of psychophysiological constructs.

## 5. Acknowledgements

This study forms part of the ReSource Project, headed by Tania Singer. Data for this project were collected between 2013 and 2016 at the former Department of Social Neuroscience at the Max Planck Institute for Human Cognitive and Brain Sciences Leipzig. We are thankful to the members of the Social Neuroscience Department involved in the ReSource Project over many years, in particular to Astrid Ackermann, Christina Bochow, Matthias Bolz and Sandra Zurborg for managing the large-scale longitudinal study, to Elisabeth Murzik, Nadine Otto, Sylvia Tydecks, and Kerstin Träger for help with recruiting and data archiving, to Henrik Grunert for technical assistance, and to Hannes Niederhausen and Torsten Kästner for data management.

## 5.2. Data availability & study materials

The present work is based on personal and sensitive physiological data that could be matched to individuals. Participants did not consent to data-sharing with parties outside the MPI CBS, such that in line with the GDPR, data cannot be made publicly available. Data are available upon reasonable request (contact via).

## 5.3. Author contributions

TS initiated, developed and secured funding for the ReSource Project. TS and VE developed and co-supervised all testing related to biomarker acquisition. Statistical analyses were performed by JS, supervised by LP and VE. SV pre-processed neuroimaging data. JS and LP drafted, and all authors contributed critically to writing the manuscript and approved its final version for submission. All authors contributed to the interpretation of the data.

