## Supplementary material for "Mapping Pathways to Neuronal Atrophy in Healthy, Mid-aged Adults: From Chronic Stress to Systemic Inflammation to Neurodegeneration?"

**to**

*Supplementary Methods A: Analysis of path associations of no interest.*

As an initial sanity check, we evaluated the significance of several paths assumed to exist between our variables but not directly related to our hypotheses. In the CT Model, age was significantly negatively associated with the latent CT factors Anterior cingulate cortex ($\beta$ =-0.51, $p < .001$), Parahippocampal Cortex ($\beta$ = -0.50, p = .012), Frontal Lobe ($\beta$ = -0.56, $p < .001$) and Temporal Lobe ($\beta$ = -0.61, $p < .001$) but not with the Entorhinal Cortex ($\beta$ = -0.27, p = .331). The latent factor Chronic Stress was significantly positively associated with hormonal status ($p \leq .001$). The latent factor Systemic Inflammation was significantly positively associated with BMI ($p \leq .001$) (see Figure 1), age ($p \leq .01$) and hormonal status ($p \leq .01$). BMI was significantly positively associated with hormonal status ($p \leq .05$) and age ($p \leq .05$). In the HCV Model, age was neither with the left nor with the right HCV significantly associated. Hormonal status was significantly positively associated with right ($\beta$ = 0.09, p = .045) but not the left HCV. The latent factor Chronic Stress was significantly negatively associated with hormonal status ($p \leq .05$) and age ($p \leq .05$). The latent factor Systemic Inflammation was significantly positively associated with BMI ($p \leq .01$) (see Figure 2), age ($p \leq .05$) and hormonal status ($p \leq .01$). BMI was significantly positively associated with hormonal status ($p \leq .05$) and age ($p \leq .05$).

*Supplementary Methods B: Handling of Heywood cases*

Heywood cases were handled by setting the product indicator variables to be equal, based on the theoretical assumption that the association between the product indicator variable and the latent interaction factor is the same for all indicator variables (Newsom, 2012) (see also Dillon et al., 1987; Kenny, 2015; Little et al., 2006). This was only done if the estimator also converged without constraining the problematic factor loadings (Chen et al., 2001). After correcting the heywood case, there were only marginally changes in model fit and coefficients, which had no impact on model significances or interpretations.

*Supplementary Methods C: Formation of latent factors of cortical thickness*

After the literature based ROI selection, we used a data driven approach to best group the following selected areas, verifying that they formed physiological valid latent factors (each separately from left and right hemispheres): Caudal Anterior Cingulate Cortex, Rostral Anterior Cingulate Cortex, Caudal Middle Frontal Cortex, Lateral Orbito-Frontal Cortex, Medial Orbito-Frontal Cortex, Parsopercularis, Parsorbitalis, Parstriangularis, Rostral Middle Frontal Cortex, Superior Frontal Cortex, Frontalpole, Paracentral cortex, precentral cortex, superior-temporal cortex, entorhinal cortex, middle-temporal cortex, temporalpole, inferiortemporal cortex, superior-temporal banks, fusiform cortex, parahippocampal cortex and transverse-temporal cortex.

To form the final five latent CT factors based on all previously selected ROI estimates, we first conducted parallel analyses (Horn, 1965) to determine the number of factors which should be considered in the factor analysis. Successively, exploratory factor analysis with all selected ROIs from both left and right hemispheres were conducted, as recommended by Westland (2010). Cortical segments with factor loadings of at least $\lambda$ = 0.4 in both hemispheres and without cross-loadings were selected. This led to the final five latent factor solution, which confirmed the topographical structure of the ROIs (see Table 1, main text). Modification indices were used to identify which of the indicator variables of CT allowed covariation of error terms with their corresponding counterpart in the other hemisphere (e.g., left hem. Frontal Superior Gyrus covarying with the right hem. Frontal Superior Gyrus). These correlations are theoretically justified through the assumption of neighbouring brain areas or hemispherical counterparts sharing additional variance that cannot be explained by the latent construct and do not significantly alter the structural or measurement parameters (Fornell, 1983; Gerbing & Anderson, 1984).

*Supplementary Methods D: Statistical pre-processing of variables*

As mentioned above, the biological variables IL-6, hs-CRP, HCC and HEC were ln-transformed to remedy their typical skewed distribution. Outliers defined as *+/- SD* = 3 were winsorized to the upper or lower boundary of 3 SDs, respectively. To account for remaining non-normality, model comparisons were evaluated with the robust Satorra-Bentler corrected $\chi^2$ (SB $\chi^2$), which can be used with non-normal distributed data (Satorra, 1988; Satorra & Bentler, 1994, 2001). For transparency, the normal $\chi^2$ as well as the SB $\chi^2$, are reported, as recommended by Curran et al.(1996). Due to their numeric distribution, observed variances of Age, BMI and HCV were at least a factor 1000 times larger than other variances, which is an obstruction for SEMs. They were thus linearly transformed by dividing them by 100 (Age and BMI) or 1000 (HCV, left and right hem.) before being added to the model.

*Supplementary Methods E: Sample Size Calculation*

As mentioned above, we assessed whether the pre-existing sample size was sufficient for the planned model using the *A-priori Sample Size Calculator for Structural Equation Models* (Soper, 2022). We expected a medium effect size of $\delta$ = .03, desired statistical power level was set to 80% and the $\alpha$-value ≤ .05. The constructed CT model consisted of 41 observed variables (incl. product indicators) and 8 latent variables, resulting in a minimum sample size of N = 108. The constructed Hippocampal model consisted of 13 observed variables (incl. product indicators) and 3 latent variables, resulting in a sample size of N = 89.

*Supplementary Table S1.* Available Samples and Reasons for Missing Values in the Raw Data.

| Variable | Available N (raw data) |
|---|---|
| Age | 332 |
| BMI | 317 |
| Sex | 332 |
| IL-6 | 315 |
| hs-CRP | 315 |
| SII | 304 |
| HCC | 210 |
| HEC | 216 |
| CT | 304 |
| HCV | 304 (282 after quality control) |

*Note:* All here reported sample size numbers refer to time-point T0 of the ReSource study. N(SII) was based on the Thrombocyte, Lymphocyte and Neutrophil count. Further details on reasons for missingness have been given elsewhere (see Engert et al., 2018 for HCC and HEC, Engert et al., 2018 and Puhlmann et al., 2019 for hs-CRP and IL-6, Valk et al. (2017) for CT, and Puhlmann, Linz et al., 2021 for HCV).

*Supplementary Table S2. Hypothesis 1 CT Model: Parameter estimates for path coefficients.*

| Regressions | Estimate | Std.Err | z-value | P(> \|z\|) | CI.low | CI.up | Std.lv | Std.all |
|---|---|---|---|---|---|---|---|---|
| ACC ~ [1] | | | | | | | | |
| Chr.Stress | 0.037 | 0.018 | 2.102 | 0.036 | 0.003 | 0.072 | 0.235 | 0.235 |
| Chr.Stress - Sys.Inf | -0.004 | 0.006 | -0.742 | 0.458 | -0.016 | 0.007 | -0.027 | -0.027 |
| FrontLobe ~ [2] | | | | | | | | |
| Chr.Stress | -0.010 | 0.013 | -0.753 | 0.451 | -0.036 | 0.016 | -0.062 | -0.062 |
| Chr.Stress - Sys.Inf | -0.002 | 0.003 | -0.521 | 0.603 | -0.008 | 0.005 | -0.010 | -0.010 |
| TempLobe ~ [3] | | | | | | | | |
| Chr.Stress | -0.009 | 0.013 | -0.700 | 0.484 | -0.035 | 0.017 | -0.056 | -0.056 |
| Chr.Stress - Sys.Inf | -0.002 | 0.003 | -0.559 | 0.576 | -0.007 | 0.004 | -0.009 | -0.009 |
| Entorhinal ~ [4] | | | | | | | | |
| Chr.Stress | 0.000 | 0.030 | 0.003 | 0.998 | -0.059 | 0.059 | 0.000 | 0.000 |
| Chr.Stress - Sys.Inf | -0.005 | 0.008 | -0.686 | 0.493 | -0.020 | 0.010 | -0.017 | -0.017 |
| Parahipp. ~ [5] | | | | | | | | |
| Chr.Stress | -0.023 | 0.028 | -0.821 | 0.412 | -0.077 | 0.032 | -0.090 | -0.090 |
| Chr.Stress - Sys.Inf | -0.005 | 0.007 | -0.747 | 0.455 | -0.018 | 0.008 | -0.019 | -0.019 |

*Note:* Parameter estimates for path coefficients of H1: [1] Direct and indirect (via Systemic Inflammation) associations of Chronic Stress with the ACC. [2] Direct and indirect (via Systemic Inflammation) associations of Chronic Stress with the Frontal Lobe. [3] Direct and indirect (via Systemic Inflammation) associations of Chronic Stress with the Temporal Lobe. [4] Direct and indirect (via Systemic Inflammation) associations of Chronic Stress with the Entorhinal Cortex. [5] Direct and indirect (via Systemic Inflammation) associations of Chronic Stress with the Parahippocampal Cortex. *Std.lv* = coefficients with only the latent variables standardized. *Std.all* = both the latent and the observed variables standardized.

*Supplementary Table S3. Hypothesis 2 CT Model: Parameter estimates for path coefficients.*

| Regressions | Estimate | Std.Err | z-value | P(> \|z\|) | CI.low | CI.up | Std.lv | Std.all |
|---|---|---|---|---|---|---|---|---|
| ACC ~ [1] | | | | | | | | |
| Sys.Inf | 1.218 | 1.140 | 1.068 | 0.286 | -1.017 | 3.453 | 0.261 | 0.236 |
| Sys.Inf x Chr.Stress | 0.600 | 0.662 | 0.906 | 0.365 | 0.697 | 1.897 | 0.144 | 0.144 |
| FrontLobe ~ [2] | | | | | | | | |
| Sys.Inf | 0.480 | 0.830 | 0.578 | 0.563 | -1.147 | 2.106 | 0.100 | 0.100 |
| Sys.Inf x Chr.Stress | -0.606 | 0.538 | -1.128 | 0.259 | -1.660 | 0.447 | -0.142 | -0.142 |
| TempLobe ~ [3] | | | | | | | | |
| Sys.Inf | 0.433 | 0.689 | 0.629 | 0.529 | -0.917 | 1.783 | 0.089 | 0.089 |
| Sys.Inf x Chr.Stress | -0.371 | 0.498 | -0.744 | 0.457 | -1.347 | 0.605 | -0.085 | -0.085 |
| Entorhinal ~ [4] | | | | | | | | |
| Sys.Inf | 1.494 | 1.775 | 0.842 | 0.400 | -1.985 | 4.974 | 0.165 | 0.165 |
| Sys.Inf x Chr.Stress | -1.750 | 1.365 | -1.282 | 0.200 | -4.425 | 0.925 | -0.217 | -0.217 |
| Parahipp. ~ [5] | | | | | | | | |
| Sys.Inf | 1.407 | 1.453 | 0.968 | 0.333 | -1.440 | 4.254 | 0.188 | 0.188 |
| Sys.Inf x Chr.Stress | 1.178 | 1.053 | 1.119 | 0.263 | -0.886 | 3.242 | 0.177 | 0.177 |

*Note:* Parameter estimates for path coefficients of H2: [1] Associations of the ACC with Systemic Inflammation and the Interaction of Systemic Inflammation and Chronic Stress on the ACC. [2] Associations of the Frontal Lobe with Systemic Inflammation and the Interaction of Systemic Inflammation and Chronic Stress on the Frontal Lobe. [3] Associations of the Temporal Lobe with Systemic Inflammation and the Interaction of Systemic Inflammation and Chronic Stress on the Temporal Lobe. [4] Associations of the Entorhinal Cortex with Systemic Inflammation and the Interaction of Systemic Inflammation and Chronic Stress on the Entorhinal Cortex. [5] Associations of the Parahippocampal Cortex with Systemic Inflammation and the Interaction of Systemic Inflammation and Chronic Stress on the Parahippocampal Cortex. *Std.lv* = coefficients with only the latent variables standardized. *Std.all* = both the latent and the observed variables standardized.

*Supplementary Table S4. Hypothesis 1 HCV Model: Parameter estimates for path coefficients.*

| Regressions | Estimate | Std.Err | z-value | P(>\|z\|) | CI.low | CI.up | Std.lv | Std.all |
|---|---|---|---|---|---|---|---|---|
| left HCV ~ [1] | | | | | | | | |
| Chr.Stress | -0.024 | 0.067 | -0.359 | 0.720 | -0.155 | 0.107 | -0.019 | -0.039 |
| Chr.Stress - Sys.Inf | 0.008 | 0.017 | 0.501 | 0.616 | -0.024 | 0.041 | 0.007 | 0.014 |
| right HCV ~ [2] | | | | | | | | |
| Chr.Stress | 0.008 | 0.063 | 0.123 | 0.902 | -0.116 | 0.131 | 0.006 | 0.013 |
| Chr.Stress - Sys.Inf | 0.019 | 0.021 | 0.910 | 0.363 | -0.022 | 0.060 | 0.015 | 0.031 |

*Note:* Parameter estimates for path coefficients of H1: [1] Direct and indirect (via Systemic Inflammation) associations of Chronic Stress with the left HCV. [2] Direct and indirect (via Systemic Inflammation) associations of Chronic Stress with the right HCV. *Std.lv* = coefficients with only the latent variables standardized. *Std.all* = both the latent and the observed variables standardized.

*Supplementary Table S5. Hypothesis 2 HCV Model: Parameter estimates for path coefficients.*

| Regressions | Estimate | Std.Err | z-value | P(>\|z\|) | CI.low | CI.up | Std.lv | Std.all |
|---|---|---|---|---|---|---|---|---|
| left HCV ∼ [1] | | | | | | | | |
| Sys.Inf | -2.311 | 4.203 | -0.550 | 0.582 | -10.548 | 5.926 | -0.054 | -0.113 |
| Sys.Inf x Chr.Stress | 0.006 | 0.504 | 0.011 | 0.991 | -0.983 | 0.994 | 0.000 | 0.001 |
| right HCV ∼ [2] | | | | | | | | |
| Sys.Inf | -5.277 | 4.684 | -1.126 | 0.260 | -14.457 | 3.904 | -0.124 | -0.261 |
| Sys.Inf x Chr.Stress | 0.740 | 0.558 | 1.326 | 0.185 | -0.354 | 1.835 | 0.041 | 0.085 |

*Note:* Parameter estimates for path coefficients of H2: [1] Associations of the left HCV with Systemic Inflammation and the Interaction of Systemic Inflammation and Chronic Stress on the left HCV. [2] Associations of the right HCV with Systemic Inflammation and the Interaction of Systemic Inflammation and Chronic Stress on the right HCV. *Std.lv* = coefficients with only the latent variables standardized. *Std.all* = both the latent and the observed variables standardized.

*Supplementary Table S6. Exploratory analysis CT Model. Parameter estimates for path coefficients.*

| Regressions | Estimate | Std.Err | z-value | P(>\|z\|) | CI.low | CI.up | Std.lv | Std.all |
|---|---|---|---|---|---|---|---|---|
| ACC ~ | | | | | | | | |
| SII | -0.032 | 0.055 | -0.576 | 0.564 | -0.140 | 0.077 | -0.281 | -0.050 |
| Chr.Stress - SII | -0.001 | 0.003 | -0.532 | 0.595 | -0.007 | 0.004 | -0.009 | -0.009 |
| FrontLobe ~ | | | | | | | | |
| SII | -0.084 | 0.050 | -1.675 | 0.094 | -0.183 | 0.014 | -0.725 | -0.129 |
| Chr.Stress - SII | -0.004 | 0.003 | -1.194 | 0.232 | -0.010 | 0.003 | -0.024 | -0.024 |
| TempLobe ~ | | | | | | | | |
| SII | -0.060 | 0.048 | -1.256 | 0.209 | -0.154 | 0.034 | -0.507 | -0.091 |
| Chr.Stress - SII | -0.003 | 0.003 | -1.050 | 0.294 | -0.008 | 0.002 | -0.017 | -0.017 |
| Entorhinal ~ | | | | | | | | |
| SII | -0.036 | 0.109 | -0.333 | 0.739 | -0.251 | 0.178 | -0.165 | -0.029 |
| Chr.Stress - SII | -0.002 | 0.005 | -0.327 | 0.744 | -0.012 | 0.008 | -0.005 | -0.005 |
| Parahipp. ~ | | | | | | | | |
| SII | -0.005 | 0.094 | -0.056 | 0.955 | -0.189 | 0.178 | -0.029 | -0.005 |
| Chr.Stress - SII | -0.000 | 0.004 | -0.056 | 0.955 | -0.009 | 0.008 | -0.001 | -0.001 |

*Note:* Parameter estimates for path coefficients for exploratory path analysis: Associations of the SII and every latent factor of CT and indirect associations of Chronic Stress via SII and every latent factor of CT. *Std.lv* = coefficients with only the latent variables standardized; *Std.all* = both the latent and the observed variables standardized.

*Supplementary Table S7.* Exploratory analysis HCV Model: Parameter estimates for path coefficients.

| Regressions | Estimate | Std.Err | z-value | P(> \|z\|) | CI.low | CI.up | Std.lv | Std.all |
|---|---|---|---|---|---|---|---|---|
| left HCV ~ | | | | | | | | |
| SII | 0.432 | 0.214 | 2.014 | 0.044 | 0.012 | 0.852 | 0.432 | 0.156 |
| Chr.Stress - SII | 0.022 | 0.014 | 1.513 | 0.130 | -0.006 | 0.050 | 0.017 | 0.035 |
| right HCV ~ | | | | | | | | |
| SII | 0.387 | 0.201 | 1.928 | 0.054 | -0.006 | 0.780 | 0.387 | 0.142 |
| Chr.Stress - SII | 0.019 | 0.013 | 1.557 | 0.119 | -0.005 | 0.044 | 0.015 | 0.032 |

*Note:* Parameter estimates for path coefficients for exploratory path analysis: Associations of the SII and left and right HCV and indirect associations of Chronic Stress via SII and left and right HCV. *Std.lv* = coefficients with only the latent variables standardized. *Std.all* = both the latent and the observed variables standardized.

*Supplementary Table S8*. Correlations of all observed variables (left hemisphere).

| | BMI | Horm. status | age | Smok. status | IL-6 | hs-CRP | SII | HCC | HEC | Fcaud ACC | Frost ACC | Parah | Entrh | caud-mid Front | Sup Front | Parac | Prec | Sup Temp | Mid Temp | Inf Temp | Stemp Banks | fus Temp | trans Temp |
|---|---|---|---|---|---|---|---|---|---|---|---|---|---|---|---|---|---|---|---|---|---|---|---|
| Horm. status | -0.22 | | | | | | | | | | | | | | | | | | | | | | |
| age | 0.2 | 0.03 | | | | | | | | | | | | | | | | | | | | | |
| Smok. status | 0.01 | 0.07 | 0.02 | | | | | | | | | | | | | | | | | | | | |
| IL-6 | 0.21 | 0.12 | 0.05 | 0 | | | | | | | | | | | | | | | | | | | |
| hs-CRP | 0.34 | 0.23 | 0.28 | 0.09 | 0.27 | | | | | | | | | | | | | | | | | | |
| SII | 0.02 | 0.07 | -0.02 | -0.05 | 0.12 | 0 | | | | | | | | | | | | | | | | | |
| HCC | 0.18 | -0.12 | 0.1 | 0.11 | -0.04 | 0.01 | 0.19 | | | | | | | | | | | | | | | | |
| HEC | 0.11 | -0.22 | -0.01 | 0.08 | -0.05 | -0.09 | 0.1 | 0.64 | | | | | | | | | | | | | | | |
| fcaudACC | 0.07 | -0.09 | -0.11 | 0.01 | 0.06 | -0.01 | -0.02 | 0.05 | 0.21 | | | | | | | | | | | | | | |
| frostACC | 0 | -0.1 | -0.3 | 0.02 | -0.08 | -0.02 | -0.04 | 0.13 | 0.13 | 0.36 | | | | | | | | | | | | | |
| Parah | 0.07 | 0 | -0.14 | 0.06 | 0.14 | 0.01 | 0.03 | 0 | -0.03 | 0.15 | 0.19 | | | | | | | | | | | | |
| Entrh | 0.05 | -0.07 | -0.07 | 0.03 | 0.02 | 0.2 | 0.07 | -0.01 | 0.04 | 0.26 | 0.14 | 0.25 | | | | | | | | | | | |
| caud-midFront | -0.02 | -0.32 | -0.36 | -0.02 | 0.13 | -0.05 | -0.15 | 0 | 0.13 | 0.28 | 0.41 | 0.18 | 0.21 | | | | | | | | | | |
| supFront | -0.07 | -0.22 | -0.41 | -0.01 | 0.06 | -0.08 | -0.19 | -0.06 | 0.02 | 0.3 | 0.42 | 0.26 | 0.16 | 0.79 | | | | | | | | | |
| Parac | -0.09 | -0.19 | -0.38 | 0.01 | 0.08 | -0.06 | -0.15 | -0.04 | 0.03 | 0.27 | 0.3 | 0.2 | 0.18 | 0.62 | 0.68 | | | | | | | | |
| Prec | -0.09 | -0.26 | -0.42 | 0.05 | -0.03 | -0.14 | -0.19 | -0.04 | 0.06 | 0.17 | 0.3 | 0.12 | 0.15 | 0.71 | 0.74 | 0.66 | | | | | | | |
| supTemp | -0.05 | -0.16 | -0.47 | 0.04 | 0.02 | -0.11 | -0.09 | -0.05 | 0.05 | 0.25 | 0.35 | 0.24 | 0.14 | 0.54 | 0.54 | 0.54 | 0.54 | | | | | | |
| midTemp | -0.16 | -0.17 | -0.47 | -0.07 | -0.02 | -0.16 | -0.16 | -0.09 | 0.05 | 0.28 | 0.41 | 0.3 | 0.24 | 0.61 | 0.64 | 0.55 | 0.63 | 0.66 | | | | | |
| infTemp | -0.09 | -0.12 | -0.22 | 0.02 | -0.15 | -0.01 | -0.15 | -0.05 | 0.07 | 0.34 | 0.41 | 0.12 | 0.27 | 0.48 | 0.49 | 0.42 | 0.45 | 0.49 | 0.63 | | | | |
| stempBanks | -0.12 | -0.05 | -0.37 | 0.11 | -0.1 | -0.14 | -0.1 | -0.04 | 0.02 | 0.17 | 0.29 | 0.14 | 0.04 | 0.39 | 0.34 | 0.33 | 0.39 | 0.6 | 0.54 | 0.42 | | | |
| fusTemp | -0.03 | 0 | -0.21 | 0.08 | 0.09 | 0.1 | -0.02 | -0.05 | 0.03 | 0.38 | 0.31 | 0.22 | 0.39 | 0.46 | 0.47 | 0.46 | 0.4 | 0.57 | 0.58 | 0.63 | 0.38 | | |
| transTemp | -0.06 | -0.03 | -0.41 | 0.04 | 0 | -0.07 | -0.03 | 0.05 | 0.07 | 0.07 | 0.14 | 0.11 | 0.03 | 0.26 | 0.34 | 0.3 | 0.36 | 0.61 | 0.35 | 0.16 | 0.32 | 0.37 | |
| HCV | 0.01 | 0.11 | 0.15 | 0.02 | 0.02 | 0.02 | 0.15 | 0.07 | -0.12 | -0.14 | -0.15 | 0.12 | 0.05 | -0.21 | -0.17 | -0.31 | -0.25 | 0.01 | -018 | -0.08 | -0.09 | -0.09 | -0.04 |

*Note:* Correlations of all preprocessed observed variables. Brain structure represented via CT and HCV in the left hemisphere. Abbrevations: hormonal status (male, female no cycle, female hormonal contraceptives, female natural cycle) (horm. Status); smoking status (smok. Status); Frontal Caudal ACC (fcaudACC); Frontal Rostral ACC (frostACC ); Parahippocampal Cortex (Parah); Entorhinal Cortex (Entrh); Frontal Caudal Middle Gyrus (caud-midFront); Frontal Superior Gyrus (supFront); Paracentral Gyrus (Parac); Precentral Gyrus (Prec); Temporal Superior Gyrus (supTemp); Temporal Middle Gyrus (midTemp ); Temporal Inferior Gyrus (infTemp); Superior Temporal Banks (stempBanks); Temporal Fusiform Gyrus (fusTemp); Transverse-temporal Gyrus (transTemp).

*Supplementary Table S9.* Correlations of all observed variables (right hemisphere).

| | BMI | Horm. status | age | Smok. status | IL-6 | hs-CRP | SII | HCC | HEC | Fcaud ACC | Frost ACC | Parah | Entrh | caud-mid Front | Sup Front | Parac | Prec | Sup Temp | Mid Temp | Inf Temp | Stemp Banks | fus Temp | trans Temp |
|---|---|---|---|---|---|---|---|---|---|---|---|---|---|---|---|---|---|---|---|---|---|---|---|
| Horm. status | -0.22 | | | | | | | | | | | | | | | | | | | | | | |
| age | 0.2 | 0.03 | | | | | | | | | | | | | | | | | | | | | |
| Smok. status | 0.01 | 0.07 | 0.02 | | | | | | | | | | | | | | | | | | | | |
| IL-6 | 0.21 | 0.12 | 0.05 | 0 | | | | | | | | | | | | | | | | | | | |
| hs-CRP | 0.34 | 0.23 | 0.28 | 0.09 | 0.27 | | | | | | | | | | | | | | | | | | |
| SII | 0.02 | 0.07 | -0.02 | -0.05 | 0.12 | 0 | | | | | | | | | | | | | | | | | |
| HCC | 0.18 | -0.12 | 0.1 | 0.11 | -0.04 | 0.01 | 0.19 | | | | | | | | | | | | | | | | |
| HEC | 0.11 | -0.22 | -0.01 | 0.08 | -0.05 | -0.09 | 0.1 | 0.64 | | | | | | | | | | | | | | | |
| fcaudACC | -0.05 | -0.09 | -0.16 | -0.02 | -0.11 | -0.12 | 0.02 | 0.07 | 0.2 | | | | | | | | | | | | | | |
| frostACC | 0.01 | -0.25 | -0.28 | 0.01 | -0.09 | -0.12 | 0.02 | 0.07 | 0.15 | 0.42 | | | | | | | | | | | | | |
| Parah | 0.01 | 0.02 | -0.14 | 0.07 | 0.06 | 0.11 | -0.06 | -0.03 | -0.1 | 0.15 | 0.16 | | | | | | | | | | | | |
| Entrh | 0.13 | -0.03 | -0.1 | 0.03 | 0.06 | 0.02 | -0.04 | 0.04 | -0.03 | 0.04 | 0.05 | 0.25 | | | | | | | | | | | |
| caud-midFront | 0.03 | -0.21 | -0.28 | -0.01 | 0.06 | -0.01 | -0.17 | -0.03 | 0.04 | 0.3 | 0.42 | 0.22 | 0.25 | | | | | | | | | | |
| supFront | -0.09 | -0.31 | -0.35 | -0.03 | -0.04 | -0.11 | -0.21 | -0.05 | -0.1 | 0.38 | 0.52 | 0.24 | 0.18 | 0.76 | | | | | | | | | |
| Parac | -0.09 | -0.21 | -0.41 | -0.08 | -0.02 | -0.14 | -0.2 | -0.08 | 0 | 0.22 | 0.29 | 0.25 | 0.13 | 0.54 | 0.62 | | | | | | | | |
| Prec | -0.08 | -0.21 | -0.4 | 0.08 | 0.03 | -0.19 | -0.17 | -0.05 | 0.06 | 0.22 | 0.44 | 0.22 | 0.18 | 0.68 | 0.7 | 0.69 | | | | | | | |
| supTemp | 0.03 | -0.23 | -0.41 | 0.07 | -0.05 | -0.13 | -0.07 | 0.02 | 0.11 | 0.15 | 0.2 | 0.29 | 0.21 | 0.43 | 0.49 | 0.46 | 0.58 | | | | | | |
| midTemp | -0.03 | -0.24 | -0.38 | -0.02 | 0 | -0.08 | -0.17 | -0.08 | -0.01 | 0.21 | 0.37 | 0.3 | 0.33 | 0.54 | 0.56 | 0.52 | 0.62 | 0.65 | | | | | |
| infTemp | 0.05 | -0.27 | -0.32 | -0.03 | -0.08 | -0.05 | -0.13 | -0.06 | -0.01 | 0.22 | 0.48 | 0.26 | 0.29 | 0.57 | 0.58 | 0.49 | 0.53 | 0.52 | 0.72 | | | | |
| stempBanks | -0.06 | -0.21 | -0.37 | -0.01 | -0.12 | -0.18 | -0.06 | -0.05 | -0.03 | 0.15 | 0.24 | 0.17 | 0.21 | 0.39 | 0.4 | 0.33 | 0.46 | 0.49 | 0.64 | 0.54 | | | |
| fusTemp | -0.02 | -0.14 | -0.28 | -0.01 | 0.03 | -0.04 | -0.06 | -0.04 | 0.06 | 0.27 | 0.31 | 0.27 | 0.41 | 0.47 | 0.52 | 0.43 | 0.48 | 0.58 | 0.66 | 0.63 | 0.4 | | |
| transTemp | -0.05 | 0.03 | -0.27 | -0.08 | -0.13 | -0.05 | 0.07 | 0.06 | 0.07 | 0.07 | 0.01 | 0.14 | 0.08 | 0.14 | 0.21 | 0.28 | 0.31 | 0.4 | 0.2 | 0.15 | 0.23 | 0.24 | |
| HCV | 0 | 0.12 | 0.12 | 0 | -0.06 | -0.04 | 0.13 | 0.09 | -0.04 | -0.03 | -0.06 | 0.02 | -0.13 | -0.14 | -0.08 | -0.17 | -0.15 | 0.04 | -0.04 | 0 | -0.07 | -0.11 | -0.04 |

*Note:* Correlations of all preprocessed observed variables. Brain structure represented via CT and HCV in the right hemisphere. Abbrevations: hormonal status (male, female no cycle, female hormonal contraceptives, female natural cycle) (horm. Status); smoking status (smok. Status); Frontal Caudal ACC (fcaudACC); Frontal Rostral ACC (frostACC ); Parahippocampal Cortex (Parah); Entorhinal Cortex (Entrh); Frontal Caudal Middle Gyrus (caud-midFront); Frontal Superior Gyrus (supFront); Paracentral Gyrus (Parac); Precentral Gyrus (Prec); Temporal Superior Gyrus (supTemp); Temporal Middle Gyrus (midTemp ); Temporal Inferior Gyrus (infTemp); Superior Temporal Banks (stempBanks); Temporal Fusiform Gyrus (fusTemp); Transverse-temporal Gyrus (transTemp).
